# Universal conditions for establish continuous cell cultures in ray-finned fishes

**DOI:** 10.1101/2023.10.17.562716

**Authors:** Adauto Lima Cardoso, Jordana Inácio Nascimento Oliveira, João Pedro Silva Climaco, Natália Bortholazzi Venturelli, Camila do Nascimento Moreira, Cesar Martins

**Affiliations:** Department of Structural and Functional Biology, Institute of Biosciences at Botucatu, Sao Paulo State University – UNESP, Botucatu, SP, Brazil

**Keywords:** Actinopterygii, biotechnology, conservation, cryopreservation, fins, invitromatics

## Abstract

Ray-finned fishes represent the most diverse vertebrate lineage and show extensive variations in physiology, ways of life and adaptations to marine and freshwater environments. Actinopterygii are largely exploited for human consumption and several species have been established as biological models. The in vitro culture of cells is fundamental for several fields of biological research, being an alternative for studies that use animals, since it mimics the cellular homogeneity of the original organisms beyond the advantage of reducing the variability of responses observed under in vivo conditions. In the case of fish cell cultures, besides being an important biomedical tool, they can offer important contributions to aquaculture and fish conservation. Hundreds of fish cell lines have been established using specific methods for each cell type and species. Here we describe an optimized protocol for obtaining cell cultures from the caudal fin of a wide range of ray-finned fishes including marine and freshwater species. The proliferative potential of these cultures makes them useful for several applications and the minimally invasive feature of this protocol makes it suitable for use in conservation plans.

## Introduction

Actinopterygii (ray-finned fishes) diverged for about 400 Mya, from the Early Devonian, and represent the most diverse fish lineage which comprises over 50% of living vertebrates and 95% of living fish species (Nelson et al. 2016). The diversity of ray-finned fishes is also reflected in extensive variations in physiology, ways of life and adaptations to marine and freshwater environments (Helfman et al. 2009). Actinopterygii are largely exploited for human consumption (Vaitla et al. 2018), and several species have been established as models to study biological processes such as development and diseases (Schartl 2014), as well as being used in toxicological studies (CarvanIII et al. 2007).

The in vitro culture of cells is fundamental for several fields of biological research, being an alternative for studies that use animals, since it mimics the cellular homogeneity of the original organisms beyond the advantage of reducing the variability of responses observed under in vivo conditions (Segeritz and Vallier 2017). In the case of fish cell cultures, besides being an important biomedical tool, they can offer important contributions to aquaculture and fish conservation (Swaminathan et al. 2021; Goswami et al. 2022). According to the Cellosaurus database (Bairoch 2018) there are 850 lines of ray-finned fishes to date, representing 196 species included in 27 orders (Fig. S1). These cell lines are derived from different organs and species, being fin, kidney, embryo, brain and gonad the most used organs (Swaminathan et al. 2021).

The caudal fin is highly variable among Actinopterygii, as a consequence of their evolutionary history and diversity of occupied environments. However, the basic structure of this fin is common among the different species, being composed of a skeleton surrounded by loose connective tissue containing nerves and blood vessels (Becerra et al. 1983; Marí-Beffa et al. 1996), and presenting high proliferative cells as the mesenchymal cells (Hou et al. 2020). Ray-finned fishes are able to regenerate injured fins by formation of the blastema, a layer of undifferentiated and proliferative cells in the wound site (Nikiforova and Golichenkov 2012). These characteristics make amputated fins and blastemas useful to development of cell cultures. Regarding the methods of cell culture, there are particularities in the protocols used for each species that do not allow their efficient use along the diversity of fishes. Therefore, here we described an optimized protocol for primary cell culture from the caudal fin of a wide range of ray-finned fishes including marine and freshwater species. The proliferative potential of these cultures makes them useful for several applications such as aquaculture, toxicology and biomedical research. The minimally invasive feature of this protocol makes it suitable for use in conservation strategies.

## Material and Methods

### Samples

Fishes were obtained from ornamental fish trade (Table 1). The animals were maintained in the fish room of the Integrative Genomic Laboratory, Sao Paulo State University, Botucatu, Brazil, under appropriate aquarium conditions and submitted to scientific procedures in accordance to the protocols of ethics in animal experimentation adopted by the Brazilian College of Experimentation Animal (COBEA) and approved by the animal ethics committee of the Institute of Biosciences-UNESP (protocol n° 486).

**Table 1.** List of fish species used for primary cell culture in the present work. Pictures of cell cultures and chromosomes are available in Supplementary Fig. S2-S21.

| Order | Species | Environment | Time for first confluence (days) | Time for confluence during passages (days) | Chromosome number (2n) | Origin of first cell culture |
| --- | --- | --- | --- | --- | --- | --- |
| Lepisosteiformes | *Lepisosteus oculatus* | Freshwater | 5 | 2 | 58 | Caudal fin – Present work |
| Osteoglossiformes | *Arapaima gigas* | Freshwater | 15 | 3 | 56 | Caudal fin – Present work |
| Cypriniformes | *Danio rerio* | Freshwater | 21 | 14 | 50 | Embryos and, pelvic fin, gill, liver and viscera of adult – Collodi et al. (1992) |
| Characiformes | *Paracheirodon axelrodi* | Freshwater | 6 | 2 | 52 | Caudal fin – Present work |
|  | *Astyanax altiparanae* | Freshwater | 3 | 2 | 50 | Muscle – Paim et al. (2018) |
|  | *Astyanax mexicanus* | Freshwater | 2 | 2 | 50 | Liver – Krishnan et al. (2022) |
| Holocentriformes | *Myripristis* sp. | Marine | 4 | 3 | 36 | Caudal fin – Present work |
| Anabantiformes | *Macropodus opercularis* | Freshwater | 3 | 2 | 46 | Caudal fin – Present work |
| Cichliformes | *Astatotilapia latifasciata* | Freshwater | 2 | 2 | 44 | Caudal fin – Present work |
|  | *Hemichromis bimaculatus* | Freshwater | 2 | 2 | 44 | Caudal fin – Present work |
|  | *Oreochromis niloticus* | Freshwater | 2 | 2 | 44 | Kidney of an interspecific hybrid – Chen et al. (1988) |
|  | *Geophagus iporangensis* | Freshwater | 2 | 2 | 48 | Caudal fin – Present work |
|  | *Pterophyllum scalare* | Freshwater | 2 | 2 | 48 | Caudal fin – Swaminathan et al. (2016) |
| Beloniformes | *Oryzias latipes* | Freshwater | 12 | 3 | 48 | Liver tumor – Mitani et al. (1980) |
|  | *Oryzias sp.* | Freshwater | 12 | 3 | 48 | Caudal fin – Present work |
| Cyprinodontiformes | *Poecilia reticulata* | Freshwater | 5 | 3 | 46 | Embryos – Li et al. (1984) |
|  | *Xiphophorus maculatus* | Freshwater | 5 | 3 | 48 | Melanoma of an interspecific hybrid<br>– Wakamatsu (1981) |
| Labriiformes | *Thalassoma lunare* | Marine | 3 | 2 | 48 | Caudal fin – Present work |
| Acanthuriformes | *Centropyge aurantonotus* | Marine | 3 | 2 | 48 | Caudal fin – Present work |
| Tetraodontiformes | *Tetraodon nigroviridis* | Freshwater | 6 | 4 | 42 | Caudal fin – Present work |

### Reagents and plastics

- 0.2% sodium hypochlorite
- 70% ethanol
- 1 × Phosphate Buffered Saline, PBS, pH 7.4
- Dulbecco’s Modified Eagle Medium/Nutrient Mixture F-12 Ham with GlutaMAX™, High Glucose, Sodium Pyruvate (DMEM/F12+GlutaMAX™, Thermo Fisher Scientific^©^, Waltham, United States of America)
- Antibiotic-Antimycotic 100 × reagent (Thermo Fisher Scientific^©^) that contains 10,000 units/mL of penicillin, 10,000 µg/mL of streptomycin, and 25 µg/mL of amphotericin B
- Collagenase Type I (1 mg/mL) (Thermo Fisher Scientific^©^)
- Fetal Bovine Serum, FBS (Cultilab^©^, Campinas, Brazil)
- TrypLE™ Express Enzyme 1 × (Thermo Fisher Scientific^©^)
- KaryoMAX™ Colcemid™ (1000 ng/mL) (Thermo Fisher Scientific^©^)
- Potassium chloride solution, KCl (0.075 M)
- Carnoy’s fixative solution (3 methanol:1 acetic acid)
- Dimethyl sulfoxide, DMSO (Sigma-Aldrich^©^)
- 25 cm^2^ cell culture flasks with filter (Sarstedt^©^, Newton, United States of America)

### Procedures

#### Biopsy collection and microbial decontamination (Fig. 1a)

Caudal fin biopsies were washed in 1 mL of 0.2% sodium hypochlorite for 10 seconds, 1 mL of 70% ethanol for 10 seconds and 1 mL of sterile 1 × PBS (pH = 7.4) for 30 seconds. Biopsies were treated in 1 mL of DMEM/F12+GlutaMAX™ supplemented with penicillin (1,000 U/mL), streptomycin (1,000 μg/mL) and amphotericin B (2.5 μg/mL) (this solution was named of treatment medium) for 2 hours. In cases such as field work, biopsies can be collected and kept in the treatment medium for days, until they are processed in the laboratory.

**Fig. 1.**
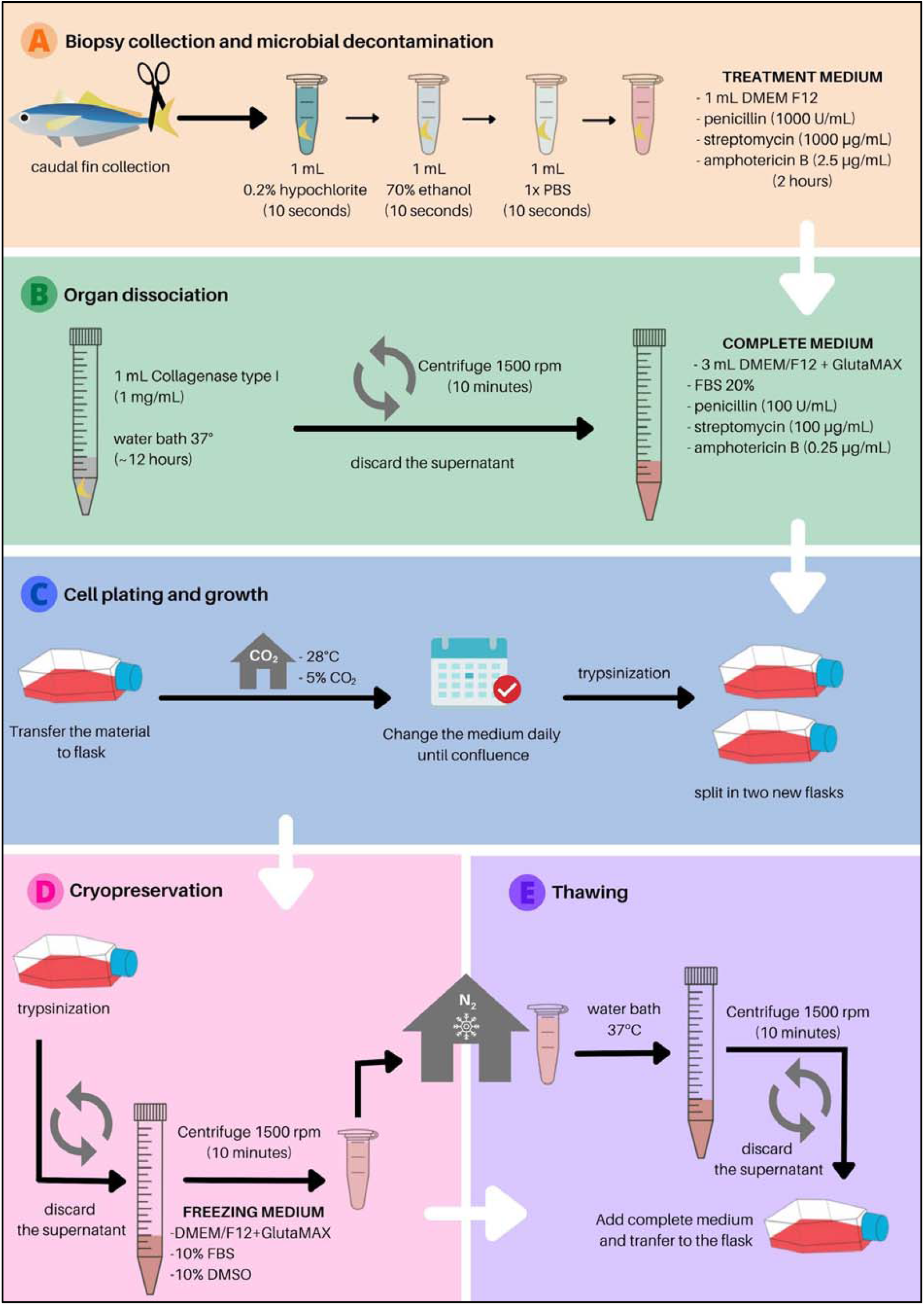
Overview of the ray-finned fish cell culture protocol. Biopsy collection and microbial decontamination (a), organ dissociation (b), cell plating and growth (c) and cryopreservation and thawing (d).

#### Organ dissociation (Fig. 1b)

Biopsies were transferred to 15 mL centrifuge tube containing 1 mL of Collagenase Type I (1 mg/mL) diluted in DMEM/F12+GlutaMAX™ and maintained in water bath for 12 hours or until complete dissociation. The material was centrifuged at 1,500 rpm for 10 minutes, the supernatant was removed and 3 mL of DMEM/F12+GlutaMAX™ medium supplemented with FBS (20%), penicillin (100 U/mL), streptomycin (100 μg/mL) and amphotericin B (0.25 μg/mL) (this solution was named of complete medium) was added and the material was homogenized by pipetting.

#### Cell plating and growth (Fig. 1c)

The homogenized material was transferred to 25 cm^2^ cell culture flasks with filter and maintained in an incubator at 28°C and 5% CO2 up 90-100% of confluence. Cells were trypsinized with TrypLE™ Express Enzyme 1 × for 3 minutes and transferred to new flasks until confluence.

#### Cryopreservation and thawing (Fig. 1d,e)

For cryopreservation, cells at 90-100% of confluence were trypsinized and transferred to DMEM/F12+GlutaMAX™ medium with 10% FBS and 10% DMSO and stored in liquid nitrogen. For revival, cells were thawed quickly in water bath at 37°C, transferred to a 15 mL centrifuge tube containing 3 mL of complete medium and centrifuged at 1,500 rpm. The supernatant was removed, the cell pellet was resuspended in 4 mL of complete medium and cells were transferred to a cell culture flask to grow.

#### Chromosome obtaining

When cells achieved the third passage, chromosome preparations were obtained by treating the cells with N-desacetyl-N-methylocolchicine contained in KaryoMAX™ Colcemid™ (10 ng/mL) for 15 minutes at 37°C. Cells were trypsinized, pelleted, hypotonized in 10 mL of KCl (0.075 M) at 37°C in 15 mL centrifuge tube immersed in water bath for 15 minutes and pre-fixed for 30 seconds by addition of 100 μL of fresh Carnoy’s fixative solution (3:1 methanol and acetic acid). Cells were pelleted, resuspended in 3 mL of fresh Carnoy’s fixative solution, which was repeated three times. The chromosome preparations were dropped onto slides and stained with Giemsa for karyotype analyzes.

## Results and Discussion

Here we presented for the first time an optimized and easily reproducible protocol for establishment of primary cultures of a wide range of Actinopterygii, including marine and freshwater species (Fig. 1, 2, 3a,b; Table 1). The great diversity of species, physiology and modes of adaptation makes the establishment of universal methods to study ray-finned fishes a challenge, which explains the development of specific methods for each cell line type. These protocols diverge in relation to the origin of cells, organ disinfection method and growth medium type. In the present work we used caudal fin for derivation of cell cultures due the presence of proliferative cells in this organ (Hou et al. 2020). Moreover, this procedure is minimally invasive due to the ability of ray-finned fishes to regenerate this organ, which is a great advantage for studies that require subsequent use of individuals or for conservation strategies, since this protocol avoids animal sacrifice. After the caudal fin collection, this organ regenerates in a few days or weeks.

**Fig. 2.**
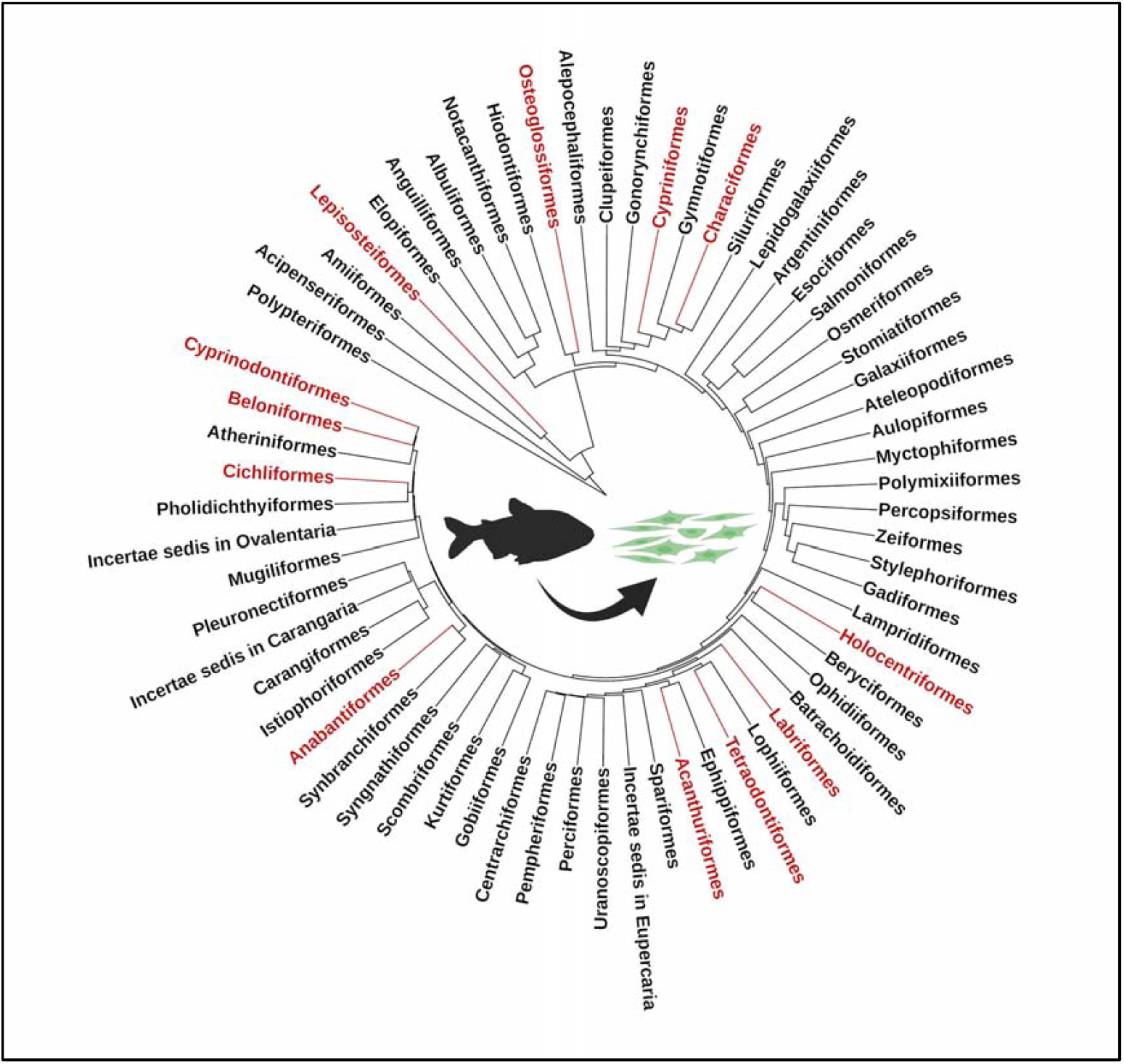
Establishment of continuous ray-finned fish cell cultures. Order-level Actinopterygii phylogeny with clades cell-cultured using the present protocol highlighted in red. Actinopterygii phylogeny was downloaded from The Fish Tree of Life (https://fishtreeoflife.org/) and edited in Interactive Tree Of Life (https://itol.embl.de/). Animal silhouettes courtesy of PhyloPic (http://www.phylopic.org).

**Fig. 3.**
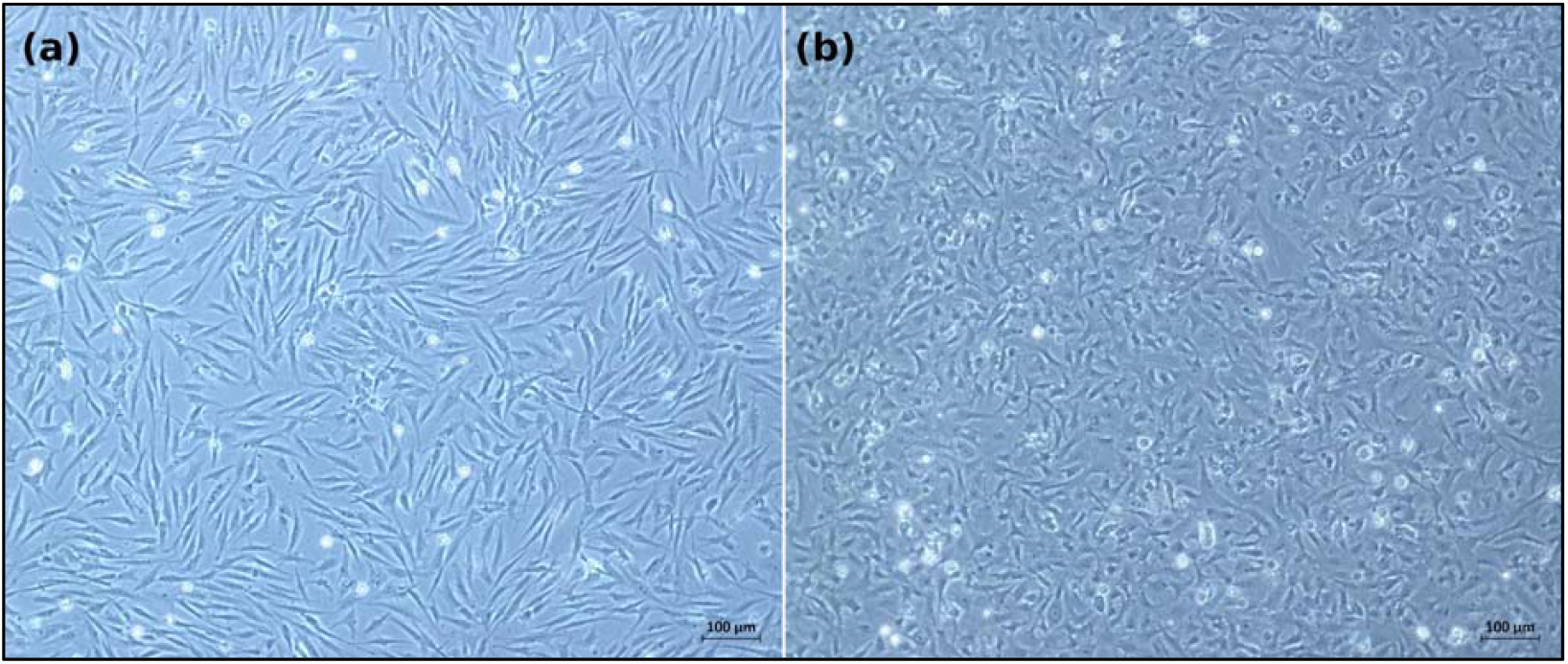
Representative primary cell cultures derived from the caudal fin of the freshwater *Arapaima gigas* (a) and the marine *Centropyge aurantonotus* (b) ray-finned fishes. Bar represents 100 μm.

### Conditions for establishment of primary cell cultures

The natural presence of microorganisms in fish organs, mainly in those exposed to external environments as skin and gills, is the first trouble to establish fish cell cultures (Akimoto et al. 2000). Microbial contamination can be eliminated by treatment with antibiotics and antimycotics. However, these agents are not always efficient to eliminate contamination from samples as their limited antimicrobial activity. Other substances such as sodium hypochlorite (NaClO) and ethanol (C2H5OH) can be used for more efficient disinfection. Due to its oxidizing action, NaClO attacks the bacterial cell wall, causing protein denaturation, while C2H5OH promotes denaturation and coagulation of microorganism proteins (Yoo 2018). Therefore, here we combined the use of NaClO, C2H5OH, antibiotics and antimycotic. To carry out a treatment with antimicrobial agents, animals can be kept in water containing these substances before organ collecting. However, this method is disadvantageous as these substances can make the subsequent use of treated individuals unfeasible, further the biological risk of excessive use of antibiotics. Moreover, this method can be expensive as it requires a large amount of drugs to obtain effective concentrations in the water. Then, in the present protocol, the caudal fin is collected and treated in a reduced volume (1 mL) of medium supplemented with high concentration of antimicrobial (Fig. 1a). The combination of these agents was effective in removing microorganisms or at least to reduce the contamination. In cases of presence of microorganisms even after the disinfection step, the elimination took place by removing the medium, followed by washing the cells with 1 × PBS and adding culture medium containing twice the concentrations indicated here.

The procedures to obtain cell cultures are categorized into enzymatic and physical methods (Jedrzejczak-Silicka 2017). In enzymatic methods, proteolytic enzymes are used for extracellular matrix disaggregation and cell dissociation. In turn, in the physical method (explant), cells start to migrate out from small pieces of original organ that are fixed in culture dishes. We conducted several tests using the explant method, but culture cells were obtained only for a few species (data not shown). On the other hand, the use of the enzyme collagenase I, that is widely used for the dissociation of many different kinds of tissues of mammals and fishes (Ryu et al. 2016), was efficient for all species tested (Fig. 1b; Table 1). Moreover, with enzyme method cell growth starts earlier (∼24h after plating) than explant (3-5 days after plating).

The culture medium is a nutritive liquid mixture composed of base medium, serum and regulatory factors, and is the most important factor in cell culture technology as it ensures cell survival, proliferation and functions (Yao and Asayama 2017; Vis et al. 2020). The determination of the type of culture medium and its supplementation is a crucial step in maintaining cell cultures. The establishment of a cell line derived from ovary of *Clarias gariepinus* required medium Leibovitz L-15 (L-15) complexly supplemented with FBS, fish muscle extract, prawn muscle extract, concanavalin A, lipopolysaccharide, glucose D, ovary extract and prawn haemolymph (Kumar et al. 2001). In turn, just L-15 medium supplemented with 15% FBS was enough to develop two cell lines derived from brains of *Epinephelus coioides* (Wen et al. 2008). The present protocol was optimized to require the least complex medium supplementation. Several types of medium and combinations of these mixtures were tested and satisfactory results were obtained using 1:1 DMEM and Ham’s F12. This combination forms a mixture that shows excellent performance, probably due to the large number of constituents in the Ham’s F12 medium and the high concentrations of the nutritional constituents of DMEM (Yao and Asayama 2017). In addition, other supplements complete the proper medium: glucose, sodium pyruvate and GlutaMAX™^©^ (Thermo Fisher Scientific). Glucose and sodium pyruvate are usually added in cell culture media as a carbon source (Yako et al. 2021). GlutaMAX™ is a substitute for L-glutamine and it minimizes toxic ammonia and improves cell viability. L-glutamine is used in cell cultures for energy production and synthesis of protein and nucleic acid. Although it is an essential nutrient, it spontaneously degrades in media, generating ammonia and pyrrolidine carboxylic acid as byproducts (Christie and Butler 1994). The mixture DMEM/F12/GlutaMAX™/high glucose/sodium pyruvate can be acquired ready-made (DMEM/F12+GlutaMAX™, Thermo Fisher Scientific^©^), as was done in the present work. The complete medium for cell maintenance and proliferation is achieved by addition of 20% FBS, penicillin (100 U/mL), streptomycin (100 μg/mL) and amphotericin B (0.25 μg/mL). This mixture was efficient for growth and maintenance of cell cultures derived from the caudal fin of twenty fish species from twelve orders, including marine and freshwater species (Table 1).

### Establishment of continuous ray-finned fish cell culture

According to the Cellosaurus database (Bairoch 2018) there are 850 cell lines of ray-finned fishes established to date, representing 196 species included in 27 orders. Perciformes, Cypriniformes and Salmoniformes are the groups with most of the cell lines available, while these three orders beside Siluriformes are the groups with most species cell-cultured (Fig. S1). Here, we developed continuous cell cultures for twenty species grouped in twelve orders (Table 1; Fig. 1, 2, 3). Twelve species were cultivated in vitro for the first time: *Lepisosteus oculatus, Arapaima gigas, Paracheirodon axelrodi, Myripristis* sp., *Macropodus opercularis, Astatotilapia latifasciata, Hemichromis bimaculatus, Melanochromis auratus, Geophagus iporangensis, Thalassoma lunare, Centropyge aurantonotus* and *Tetraodon nigroviridis*. Cell culture of a species of Holocentriformes (*Myripristis* sp.) was also performed for the first time here. Our results show that the present protocol is efficient to establish continuous cell cultures for several fish lineages that started to diverge about 400 Mya. In this manner, we suggest that this protocol should be expanded and applied even to groups of Actinopterygii and other tissue/organs that were not tested here, contributing to the expansion of the field known as invitromatics (Bols et al. 2017). The great proliferative ability of these cultures indicates that they can acquire immortality, passing to be considered cell lines. Cell cultures obtained here showed conserved diploid chromosome numbers in comparison with in vivo descriptions (Fig. 4; Fig. S2-S21), which suggest these cultures have stable genomes. Both primary cultures and cell lines have potential to be used in several fields, such as cytogenetics, genomics, pathology, immunology, toxicology, cellular agriculture and conservation (Swaminathan et al. 2021; Goswami et al. 2022).

**Fig. 4.**
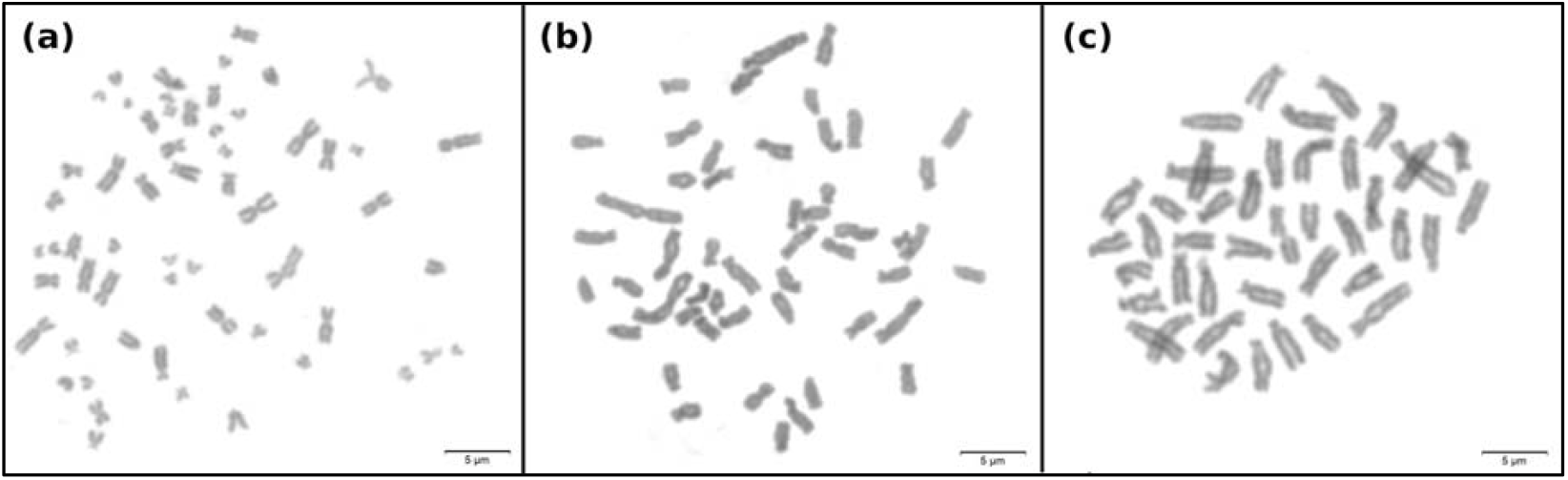
Representative mitotic metaphase chromosomes obtained from primary cell cultures derived from the caudal fin of *Lepisosteus oculatus* showing 2n = 58 (a), *Astyanax mexicanus* showing 2n = 50 (b) and *Astatotilapia latifasciata* (c) showing 2n = 44. Bar represents 5 μm.

## Conclusions

This study allowed for the first time the optimization of a protocol for use in a very wide range of ray-finned fishes, allowing the establishment of cell cultures in a standardized way, contributing to the invitromatics of Actinopterygii. The derivation of cells by this protocol has application in several fields of research and this method has the potential to be expanded to species not yet tested, as well as to other organs.

## Acknowledgments

This work was supported by grants from São Paulo Research Foundation (FAPESP) (grant numbers 2015/16661-1; 2017/07484-4; 2017/25193-7; 2018/09553-6; 2018/25350-8; 2022/02544-7) and the National Council for Scientific and Technological Development (CNPq) to C.M.

## Author Contribution

ALC designed and coordinated the study, performed all the experiments and wrote the manuscript. JIN, JPSC and NBV contribute to cell growth and maintenance. CNM contributed to karyotyping procedures, cell growth and maintenance. CM participated in design and coordination of the study. All the authors read and approved the final version of the manuscript.

## Data Availability

All data relevant to the study are included in the article. Additional data are accessible in the electronic Supplementary Material.

## Ethics approval

Animals were maintained and submitted to scientific procedures in accordance with the protocols of ethics in animal experimentation adopted by the Brazilian College of Experimentation Animal (COBEA) and approved by the animal ethics committee of the Institute of Biosciences-UNESP (protocol n° 1833090223).

## Consent to participate

All the authors participated in analyses and interpretation of data and read and approved the final manuscript.

## Consent for publication

All the authors approved the final manuscript for publication.

