## Supplemental Figures for "Universal conditions for establish continuous cell cultures in ray-finned fishes"

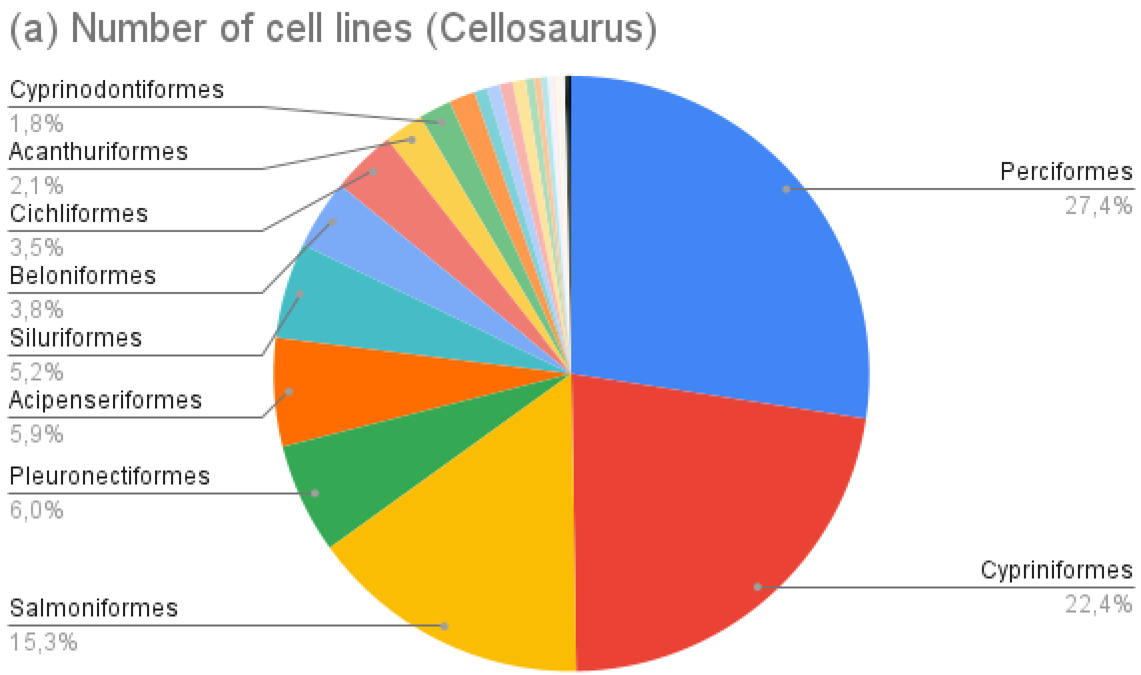


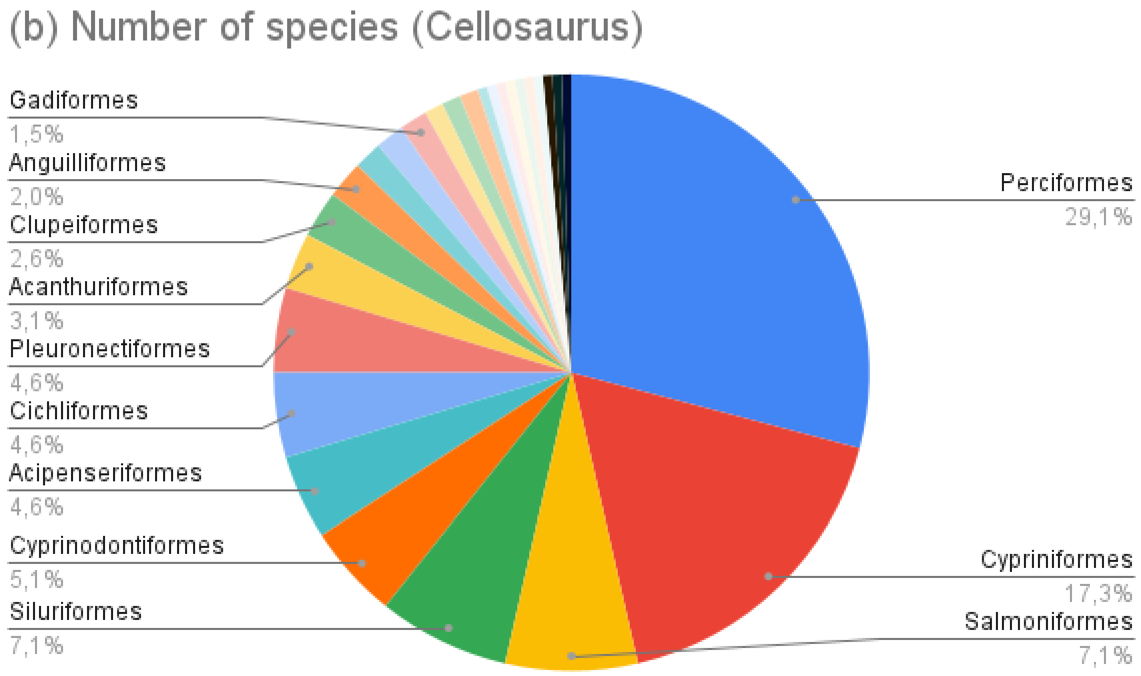


**Supplementary Fig. S1** Collection of ray-finned fish cell lines available in Cellosaurus organized in level of order. Number of cell lines (a). Number of species with cell lines established.

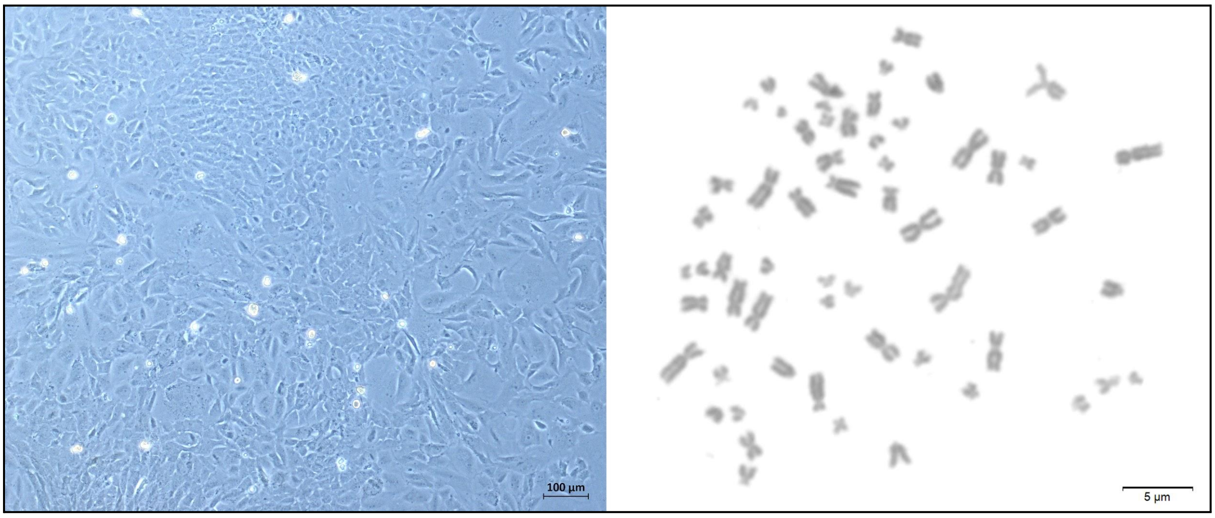


**Supplementary Fig. S2** *Lepisosteus oculatus* primary cell culture derived from caudal fin at third passage (left) and metaphase showing 2n = 58 chromosomes (right) as previously observed (Braasch et al., 2016).

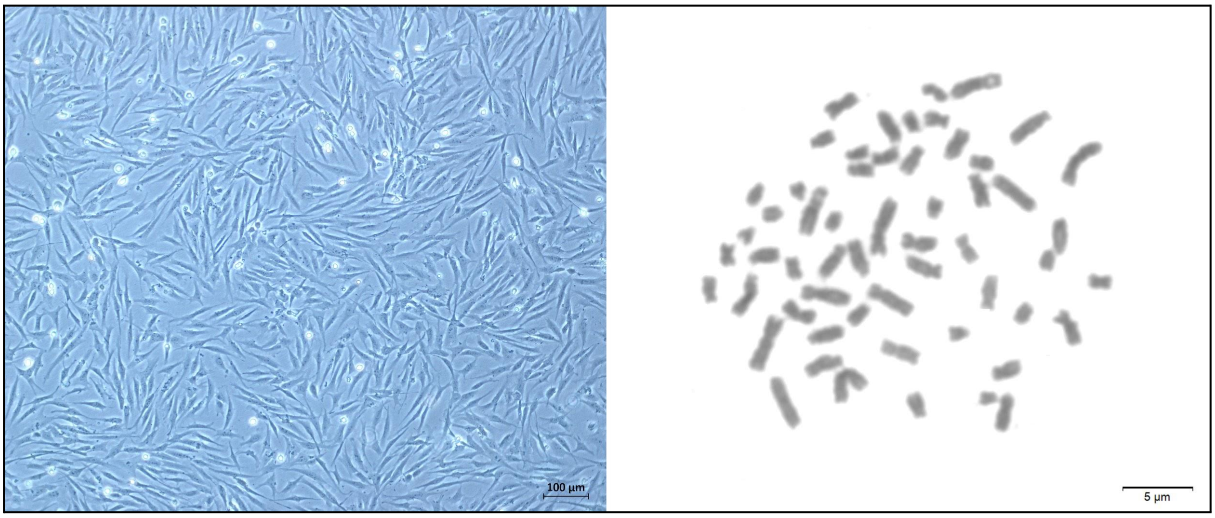


**Supplementary Fig. S3** *Arapaima gigas* primary cell culture derived from caudal fin at third passage (left) and metaphase showing 2n = 56 chromosomes (right) as previously observed (Marques et al., 2006).

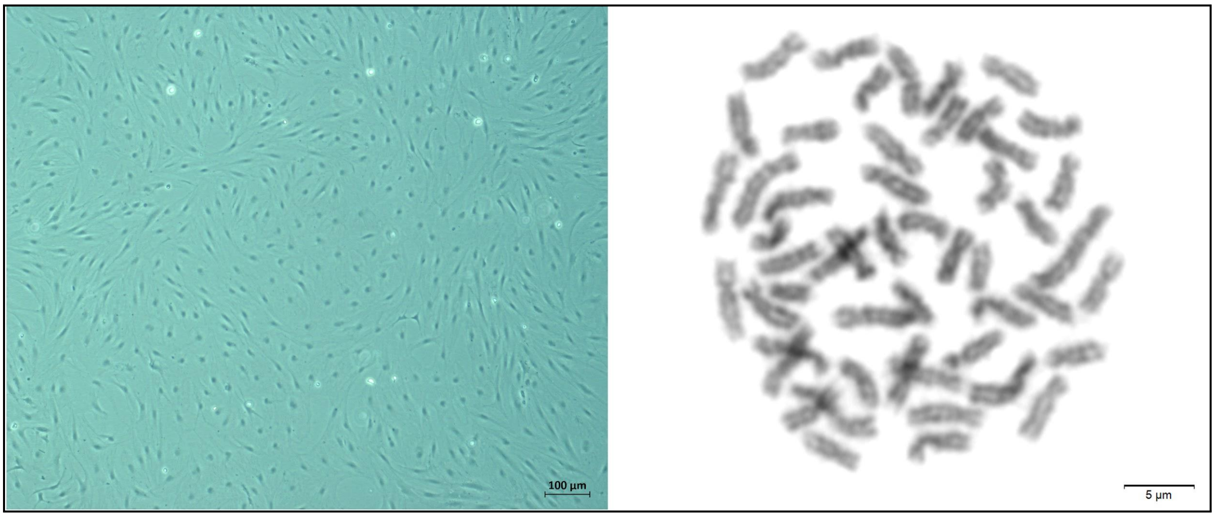


**Supplementary Fig. S4** *Danio rerio* primary cell culture derived from caudal fin at third passage (left) and metaphase showing 2n = 50 chromosomes (right) as previously observed (Sola & Gornung, 2001).

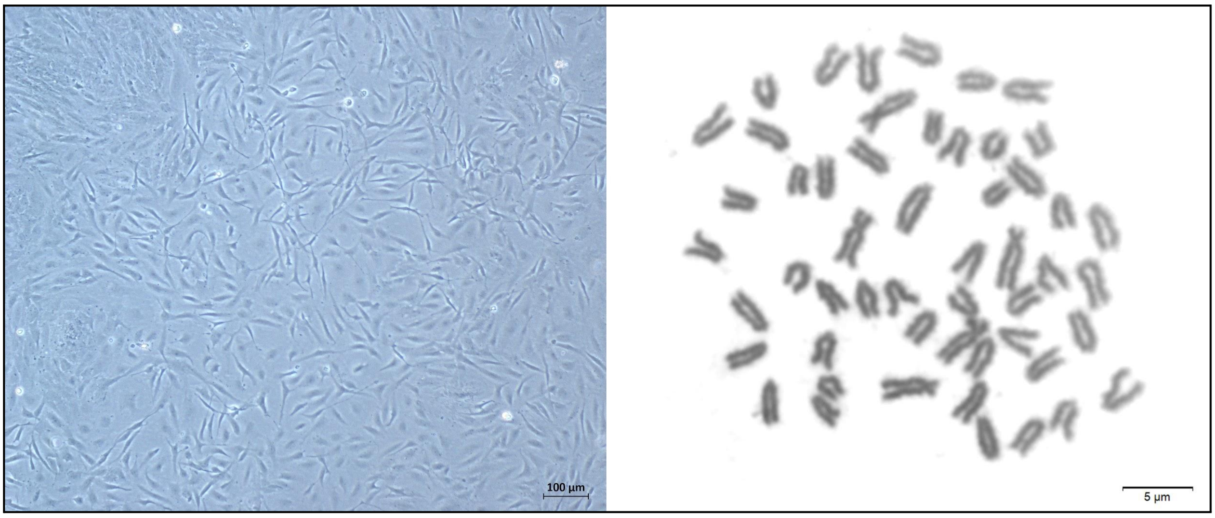


**Supplementary Fig. S5** *Paracheirodon axelrodi* primary cell culture derived from caudal fin at third passage (left) and metaphase showing 2n = 52 chromosomes (right) as previously observed (Scheel & Christensen, 1972).

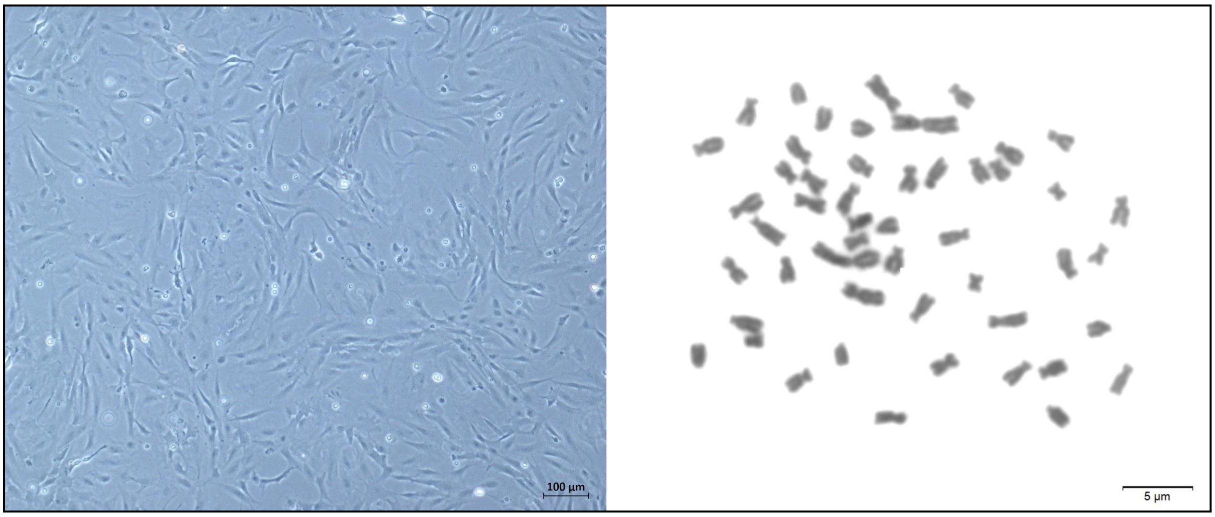


**Supplementary Fig. S6** *Astyanax altiparanae* primary cell culture derived from caudal fin at third passage (left) and metaphase showing 2n = 50 chromosomes (right) as previously observed (Ferreira Neto, 2009).

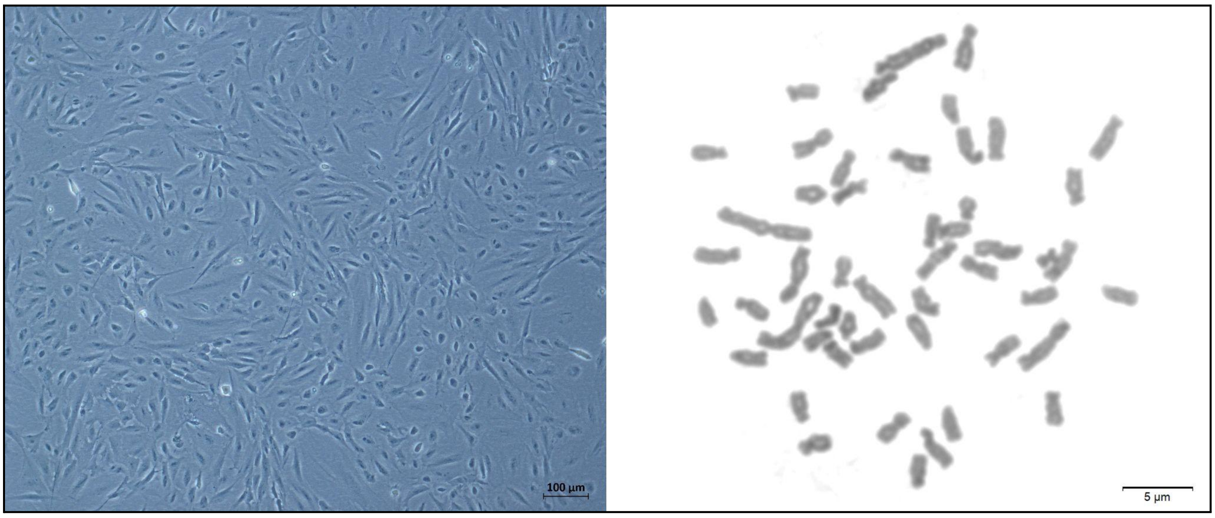


**Supplementary Fig. S7** *Astyanax mexicanus* primary cell culture derived from caudal fin at third passage (left) and metaphase showing 2n = 50 chromosomes (right) as previously observed (Scheel, 1972).

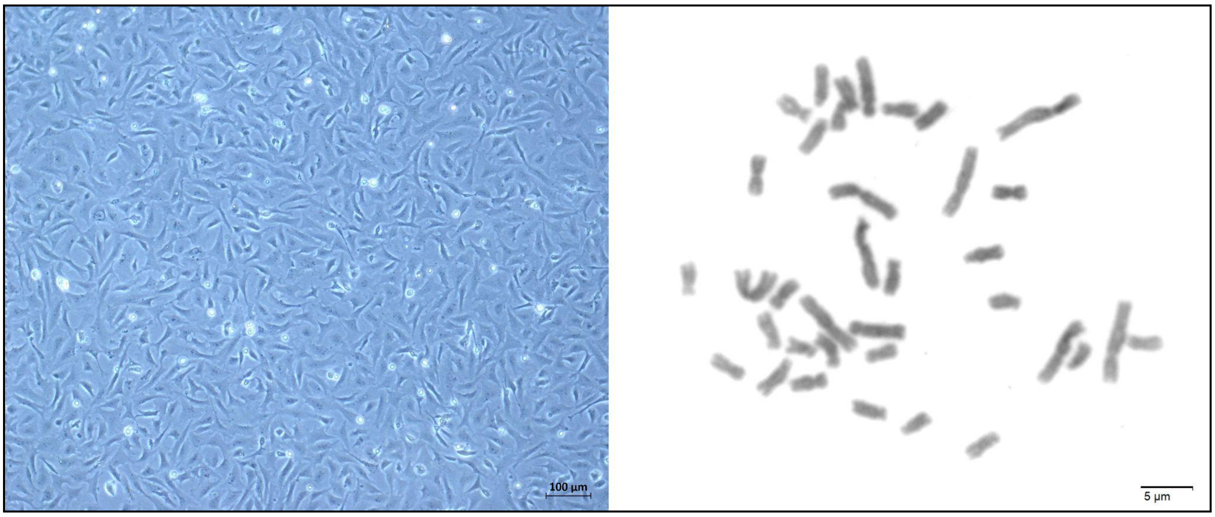


**Supplementary Fig. S8** *Myripristis* sp. primary cell culture derived from caudal fin at third passage (left) and metaphase showing 2n = 36 chromosomes (right).

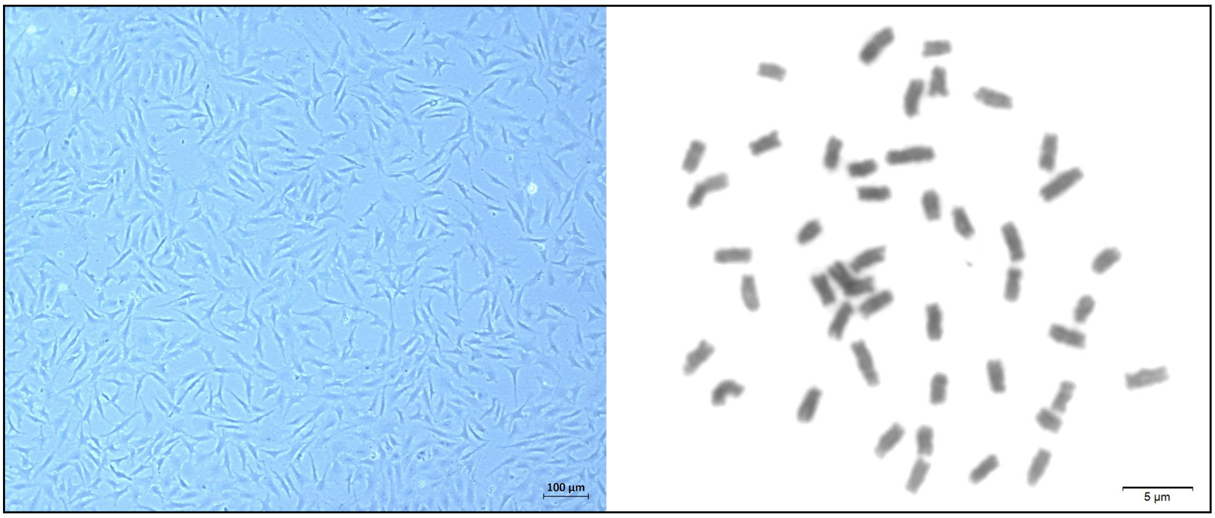


**Supplementary Fig. S9** *Macropodus opercularis* primary cell culture derived from caudal fin at third passage (left) and metaphase showing 2n = 46 chromosomes (right) as previously observed (Abe, 1975).

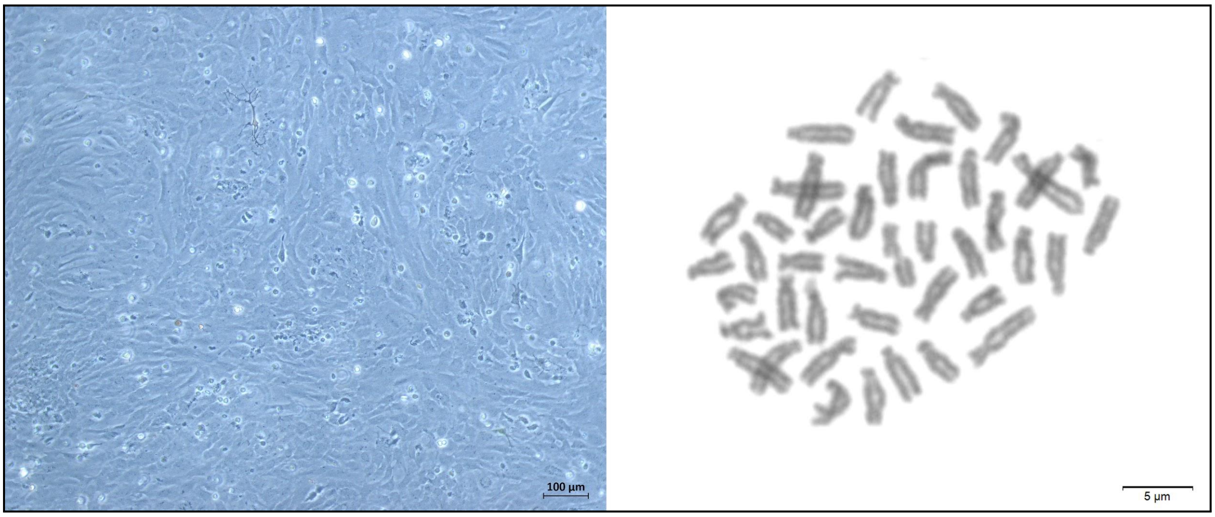


**Supplementary Fig. S10** *Astatotilapia latifasciata* primary cell culture derived from caudal fin at third passage (left) and metaphase showing 2n = 44 chromosomes (right) as previously observed (Poletto el al., 2010a).

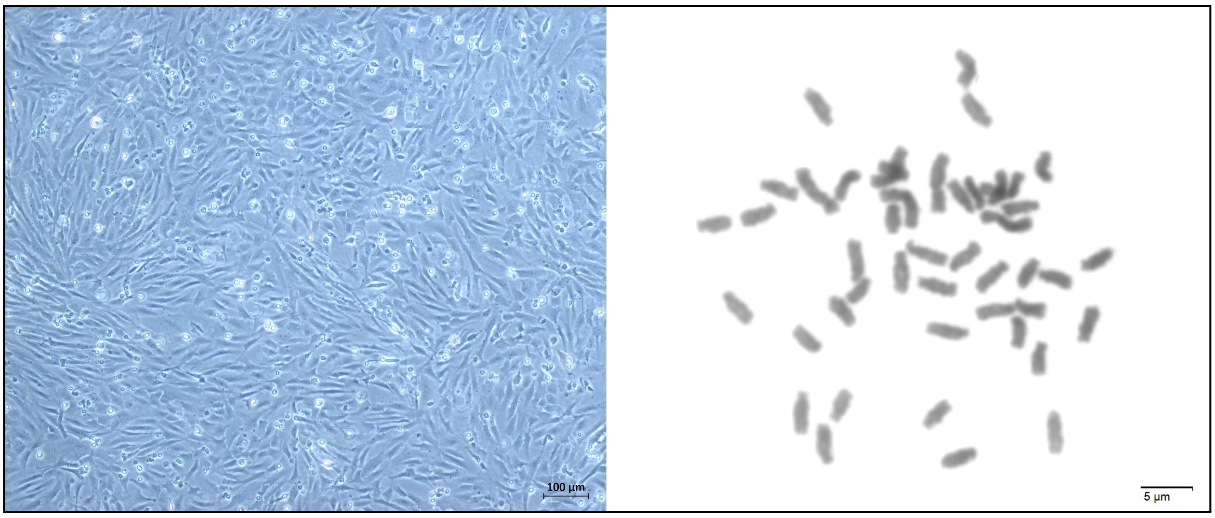


**Supplementary Fig. S11** *Hemichromis bimaculatus* primary cell culture derived from caudal fin at third passage (left) and metaphase showing 2n = 44 chromosomes (right) as previously observed (Poletto et al., 2010b) .

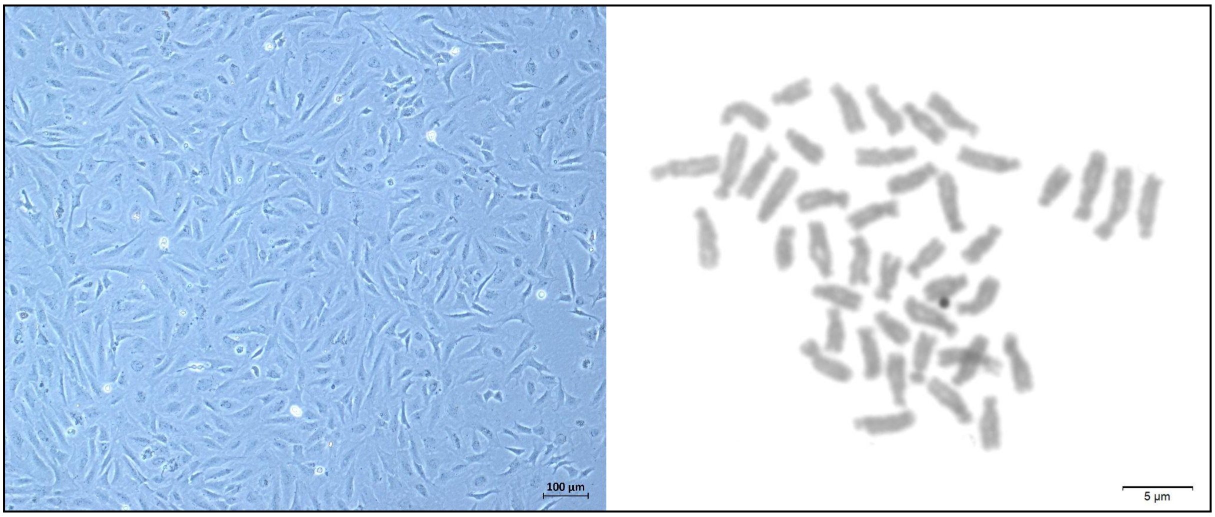


**Supplementary Fig. S12** *Oreochromis niloticus* primary cell culture derived from caudal fin at third passage (left) and metaphase showing 2n = 44 chromosomes (right) as previously observed (Chervinski, 1964).

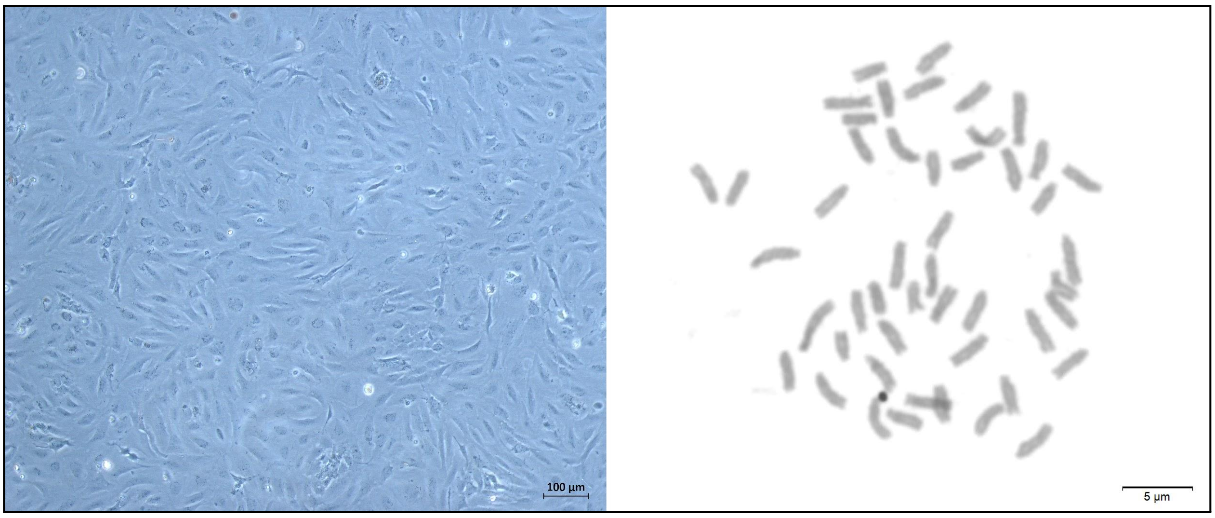


**Supplementary Fig. S13** *Geophagus iporangensis* primary cell culture derived from caudal fin at third passage (left) and metaphase showing 2n = 48 chromosomes (right).

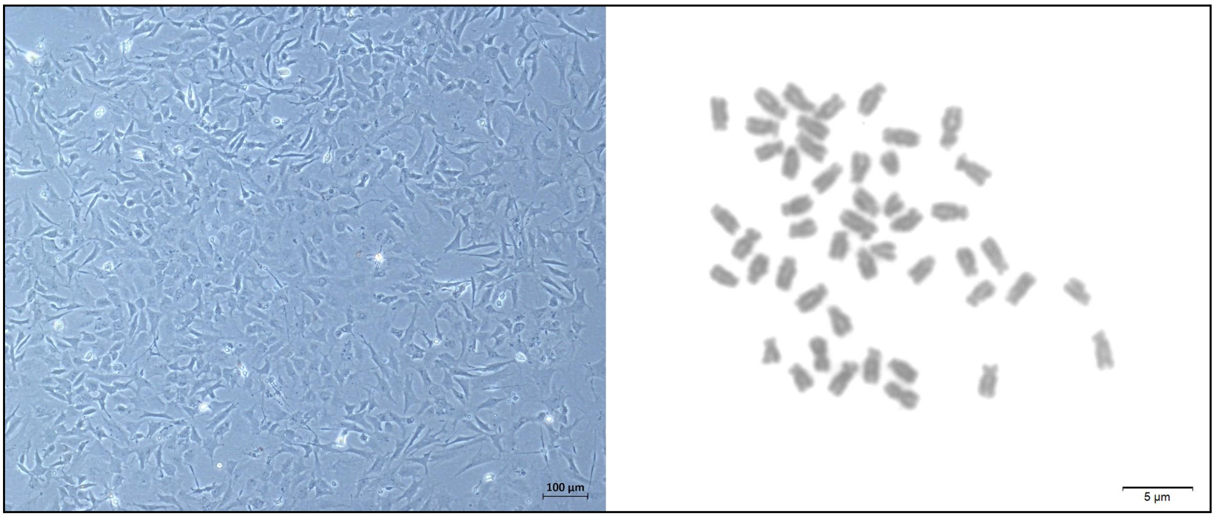


**Supplementary Fig. S14** *Pterophyllum scalare* primary cell culture derived from caudal fin at third passage (left) and metaphase showing 2n = 48 chromosomes (right) as previously observed (Thompson, 1979).

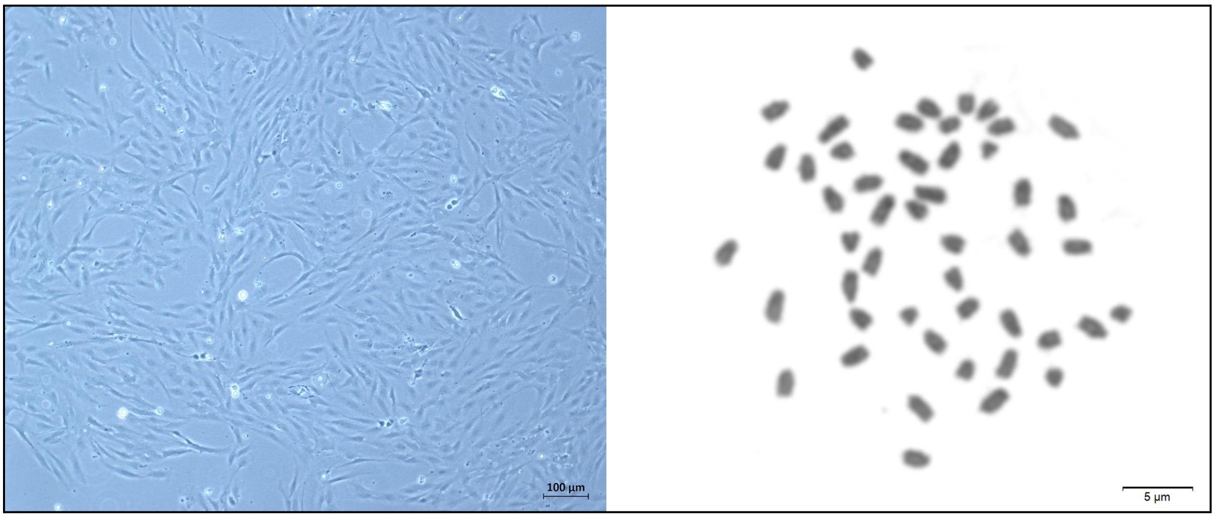


**Supplementary Fig. S15** *Oryzias latipes* primary cell culture derived from caudal fin at third passage (left) and metaphase showing 2n = 48 chromosomes (right) as previously observed (Ojima & Hitotsumachi, 1969).

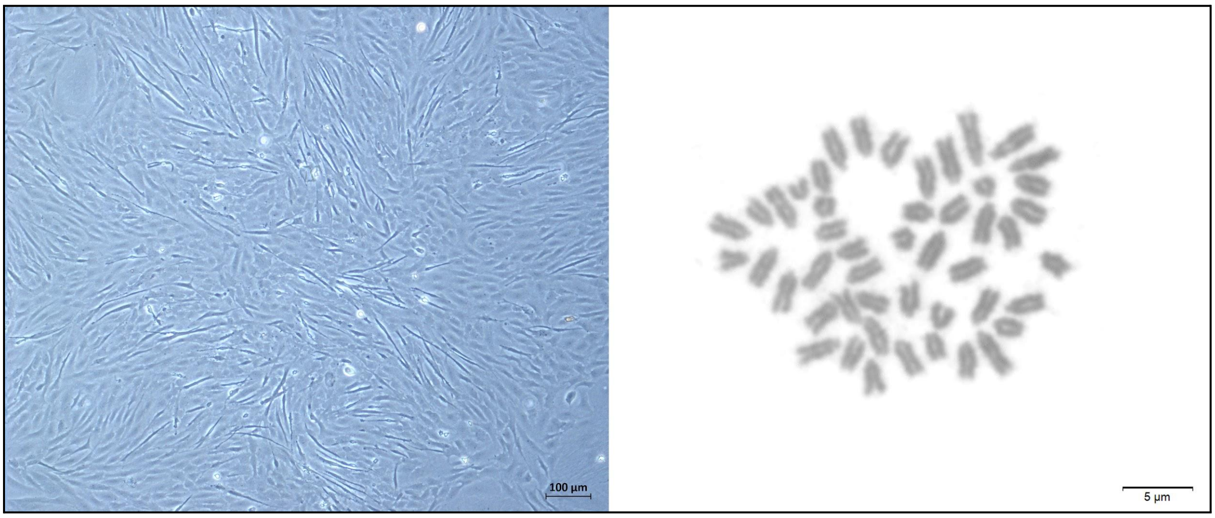


**Supplementary Fig. S16** *Oryzias* sp. primary cell culture derived from caudal fin at third passage (left) and metaphase showing 2n = 48 chromosomes (right).

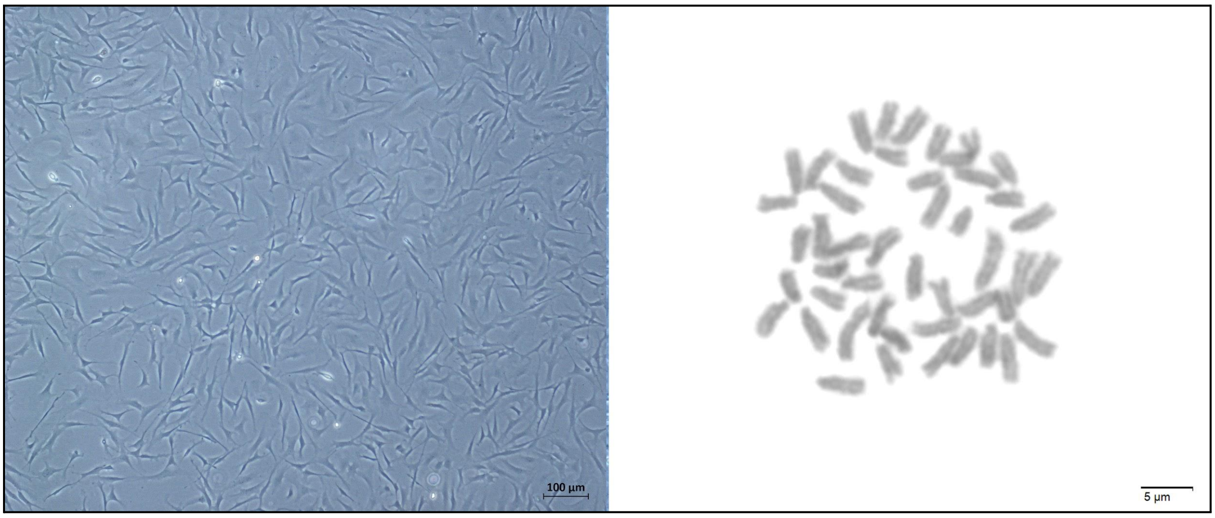


**Supplementary Fig. S17** *Poecilia reticulata* primary cell culture derived from caudal fin at third passage (left) and metaphase showing 2n = 46 chromosomes (right) as previously observed (Winge, 1922).

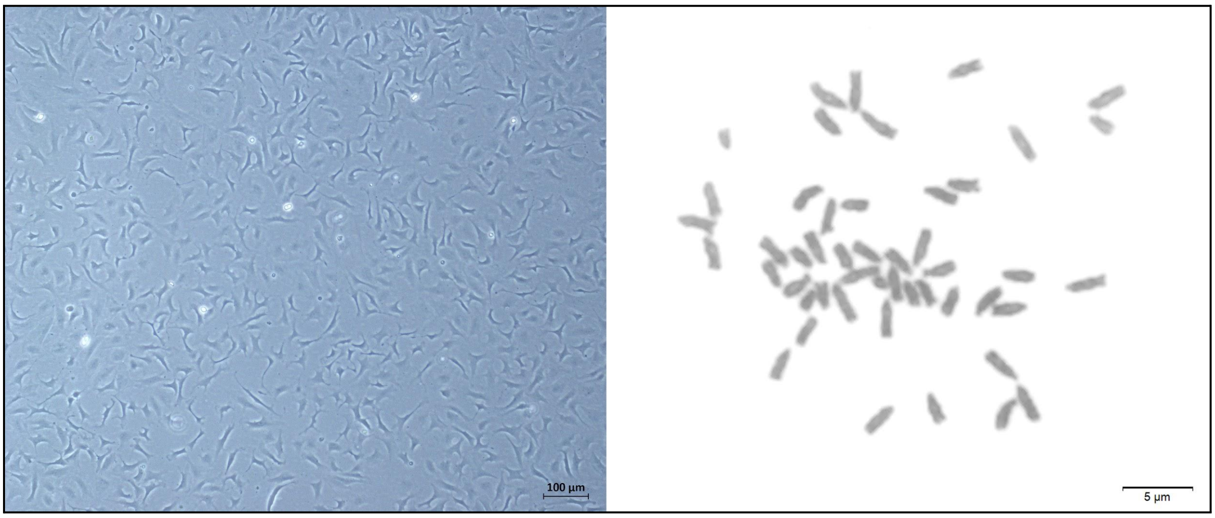


**Supplementary Fig. S18** *Xiphophorus maculatus* primary cell culture derived from caudal fin at third passage (left) and metaphase showing 2n = 48 chromosomes (right) as previously observed (Nanda et al., 1993).

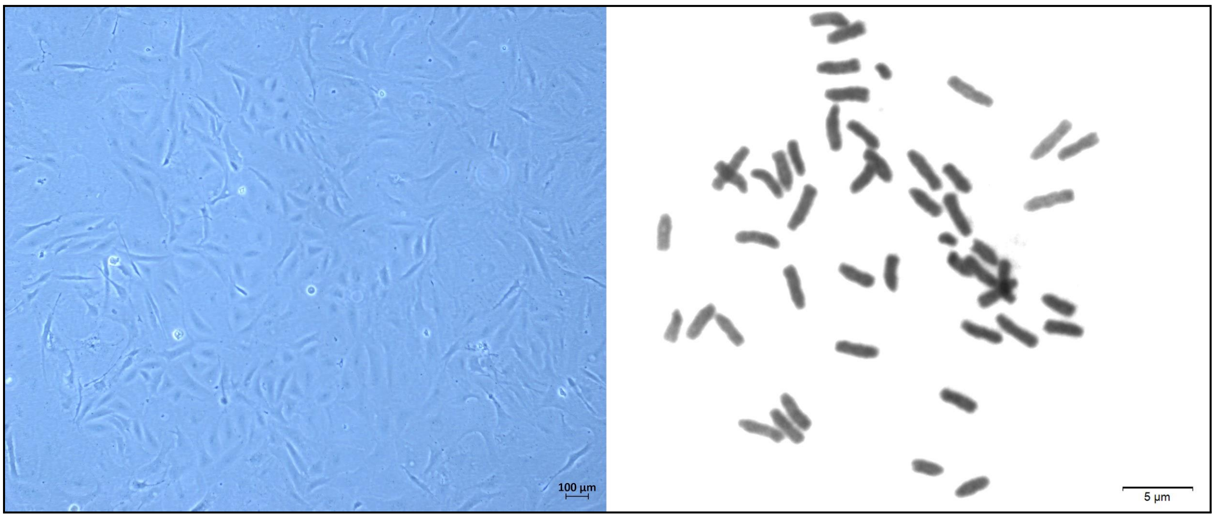


**Supplementary Fig. S19** *Thalassoma lunare* primary cell culture derived from caudal fin at third passage (left) and metaphase showing 2n = 48 chromosomes (right) as previously observed (Ojima & Kashiwagi, 1979).

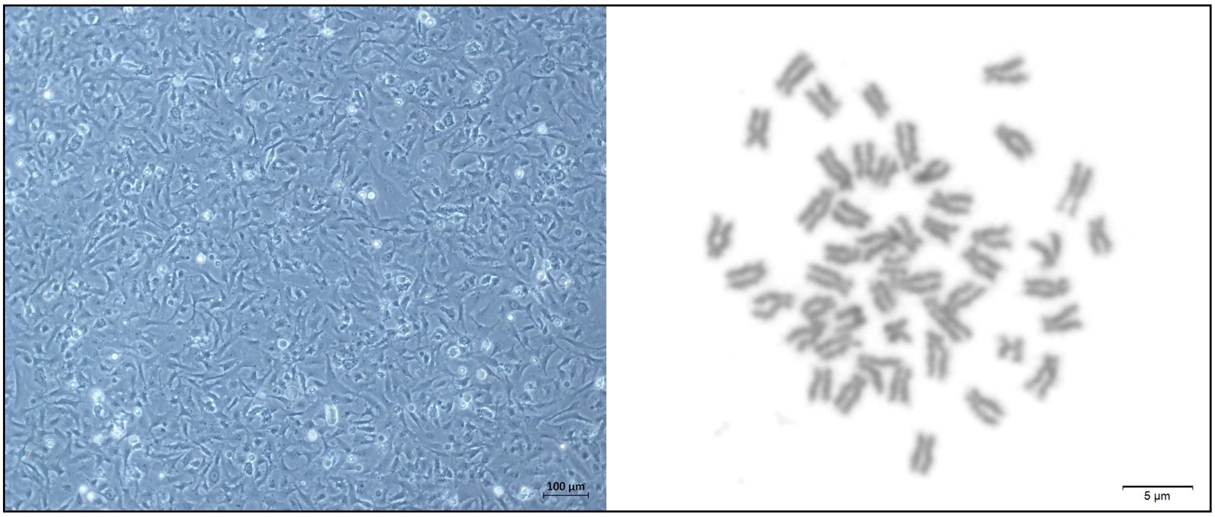


**Supplementary Fig. S20** *Centropyge aurantonotus* primary cell culture derived from caudal fin at third passage (left) and metaphase showing 2n = 48 chromosomes (right) as previously observed (Affonso & Galleti-Jr, 2005).

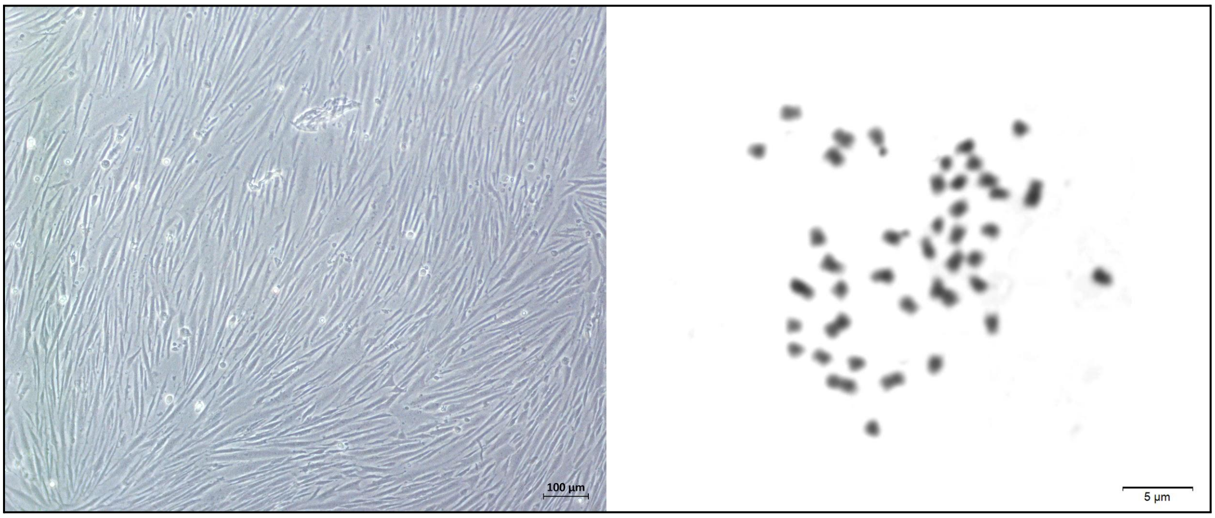


**Supplementary Fig. S21** *Tetraodon nigroviridis* primary cell culture derived from caudal fin at third passage (left) and metaphase showing 2n = 42 chromosomes (right) as previously observed (Fischer et al., 2000).
